## Supplementary figures and images for "DePARylation is critical for S phase progression and cell survival"

### Figure 1-Figure Supplement 1

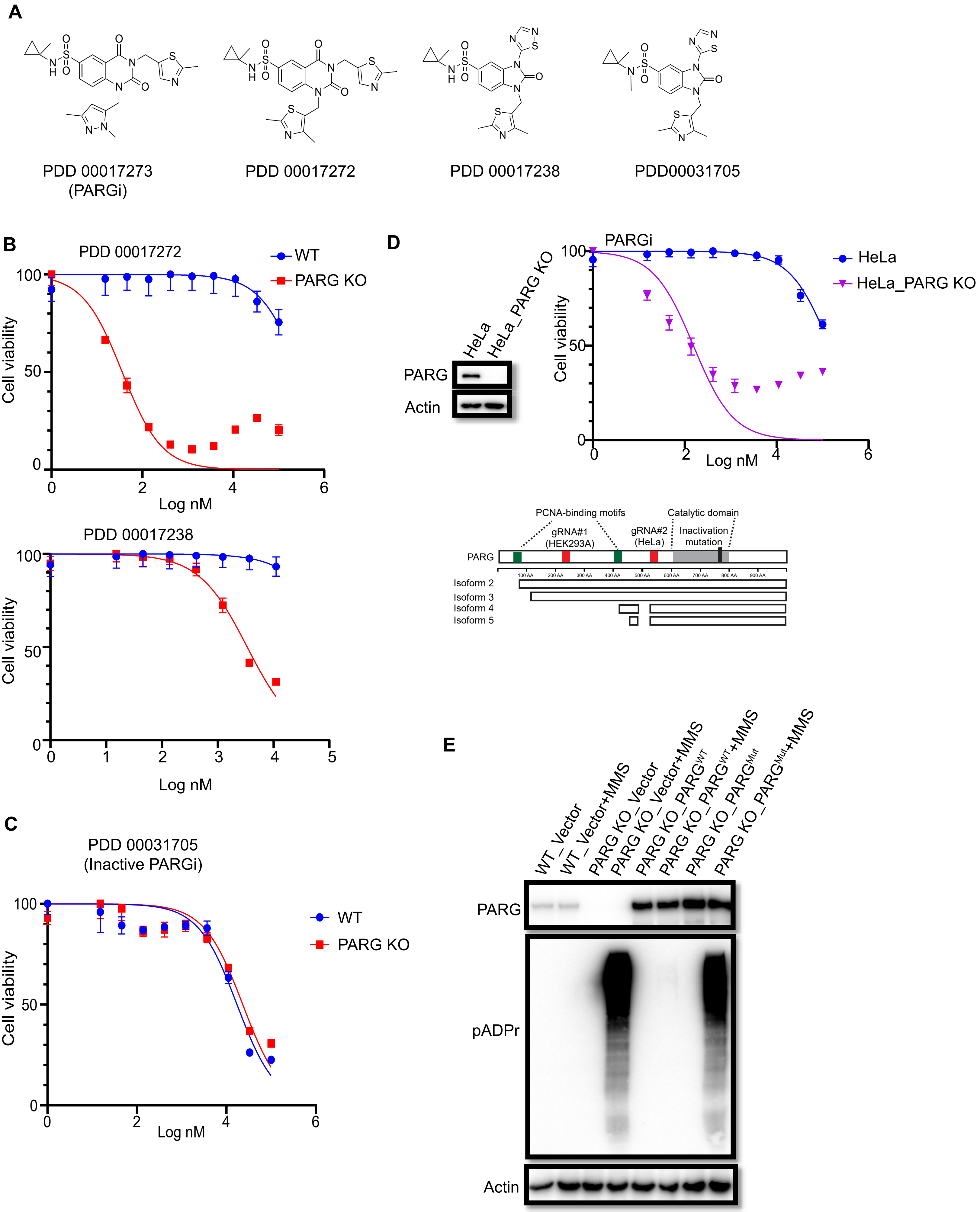

### Figure 2-Figure Supplement 1

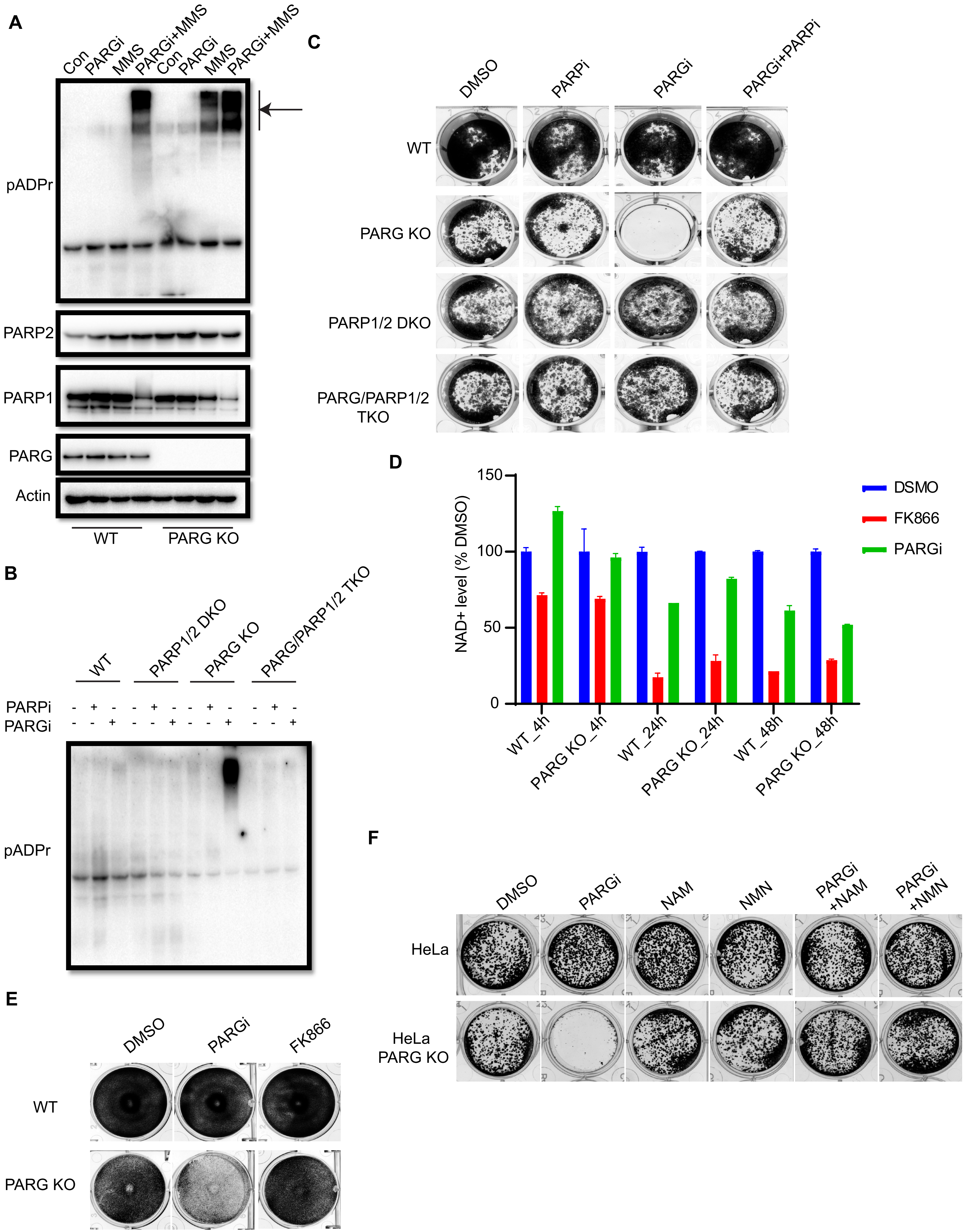

### Figure 3-Figure Supplement 1

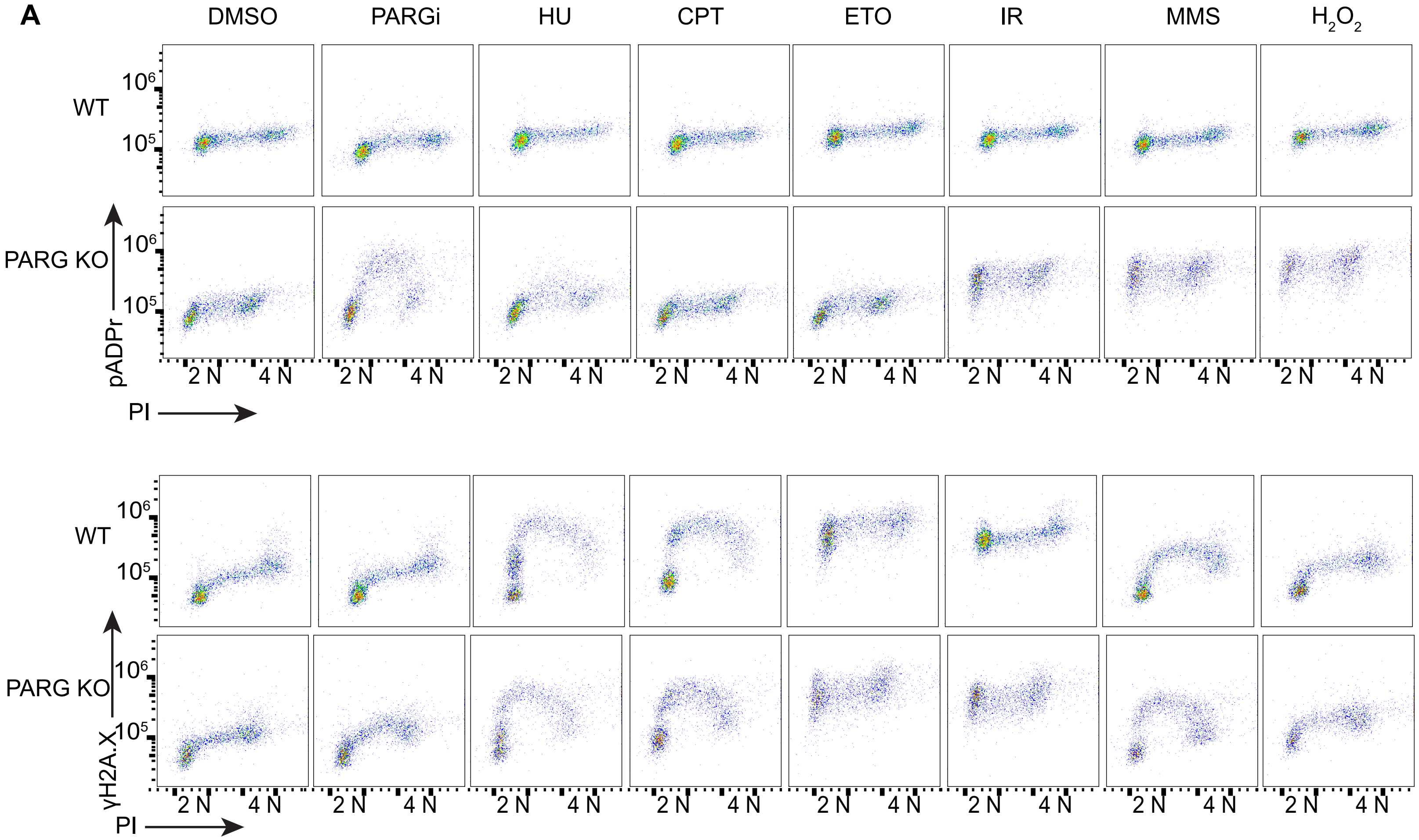

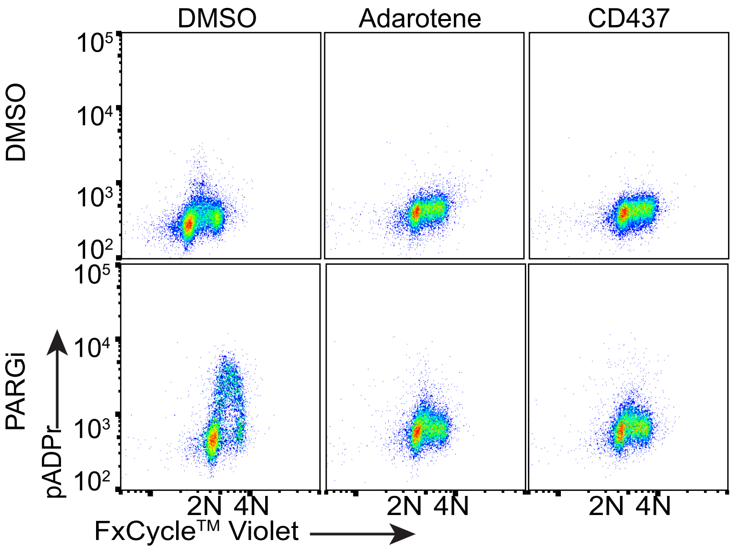

### Figure 4-Figure Supplement 1

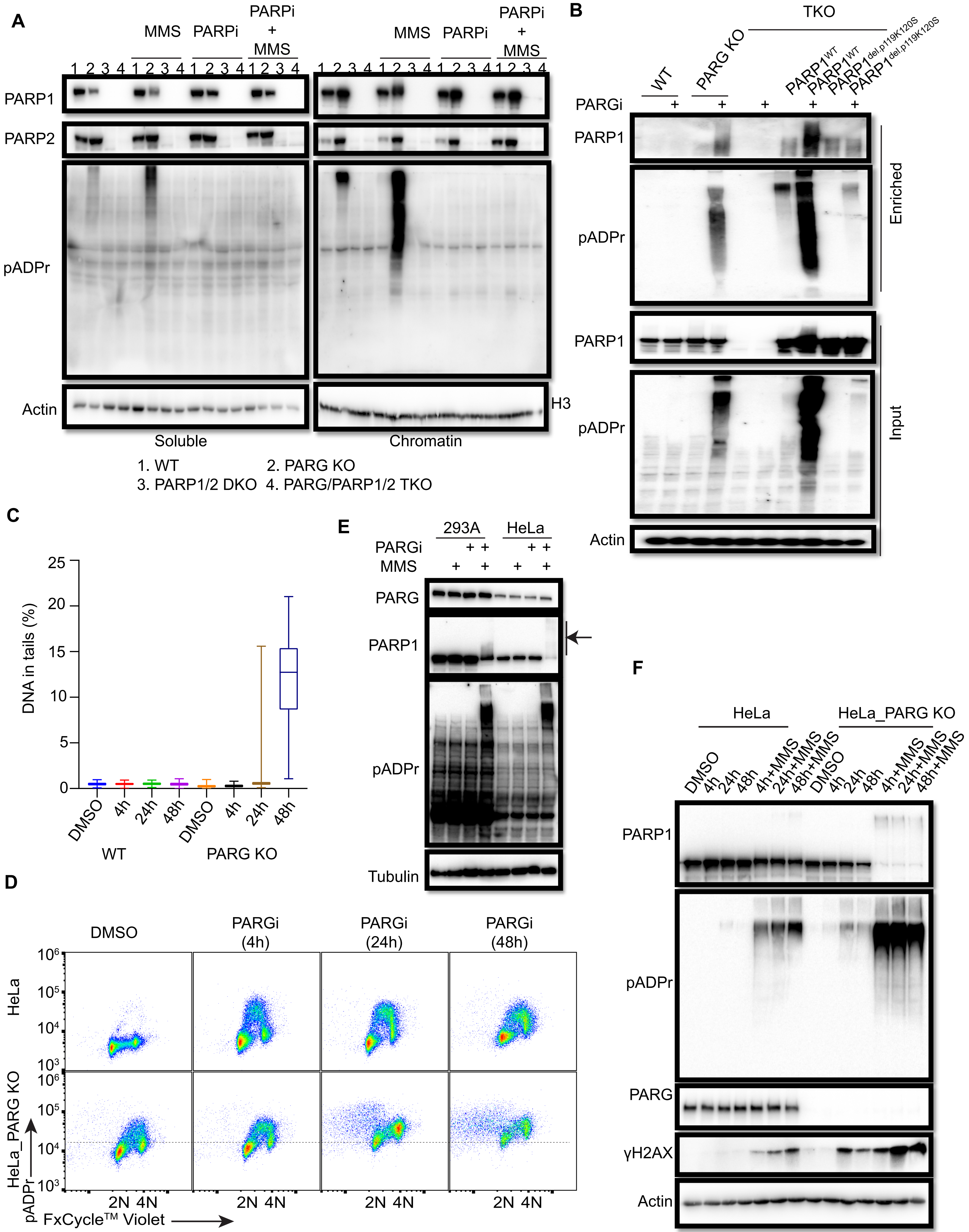

### Figure 5-Figure Supplement 1

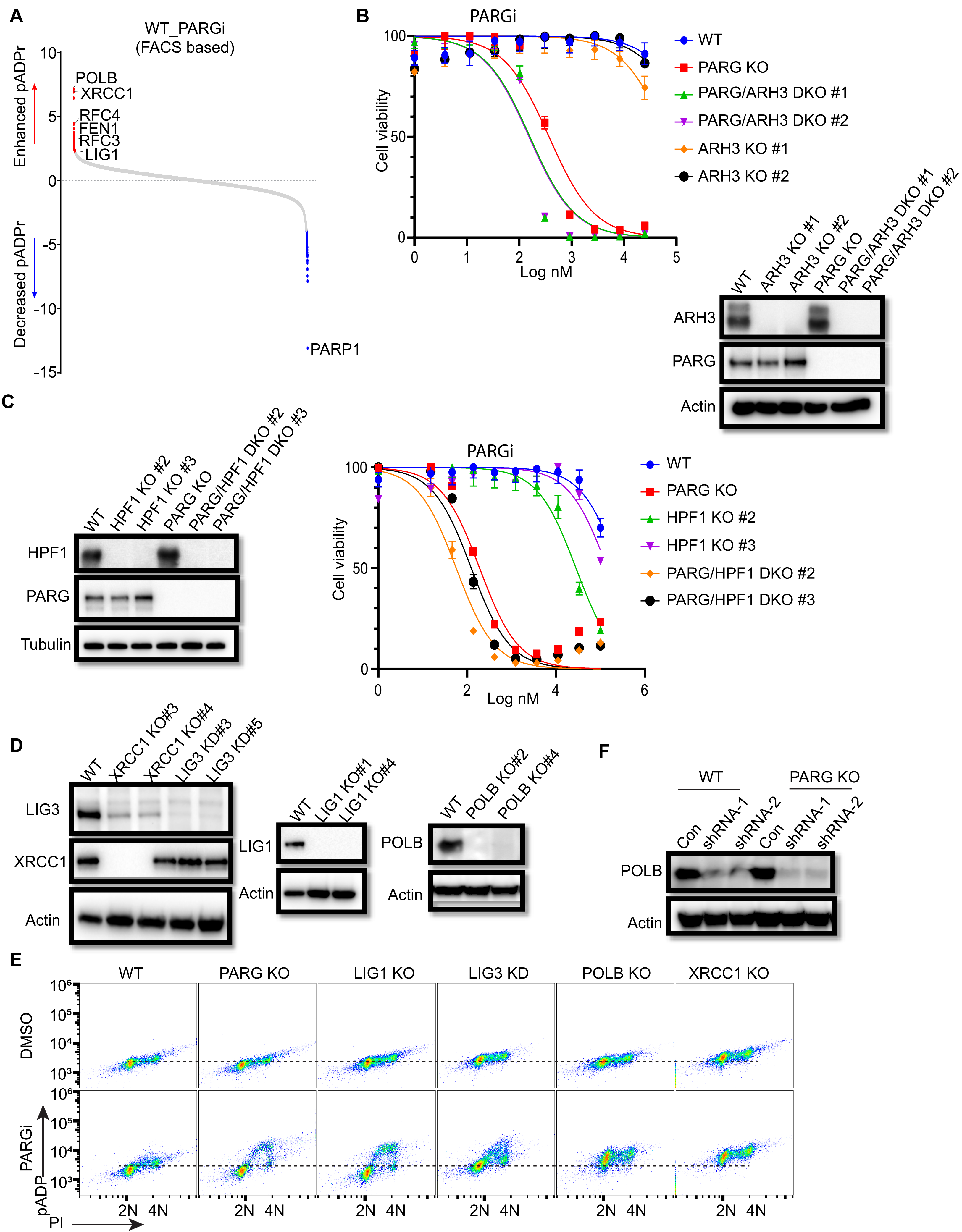

### Figure 6-Figure Supplement 1

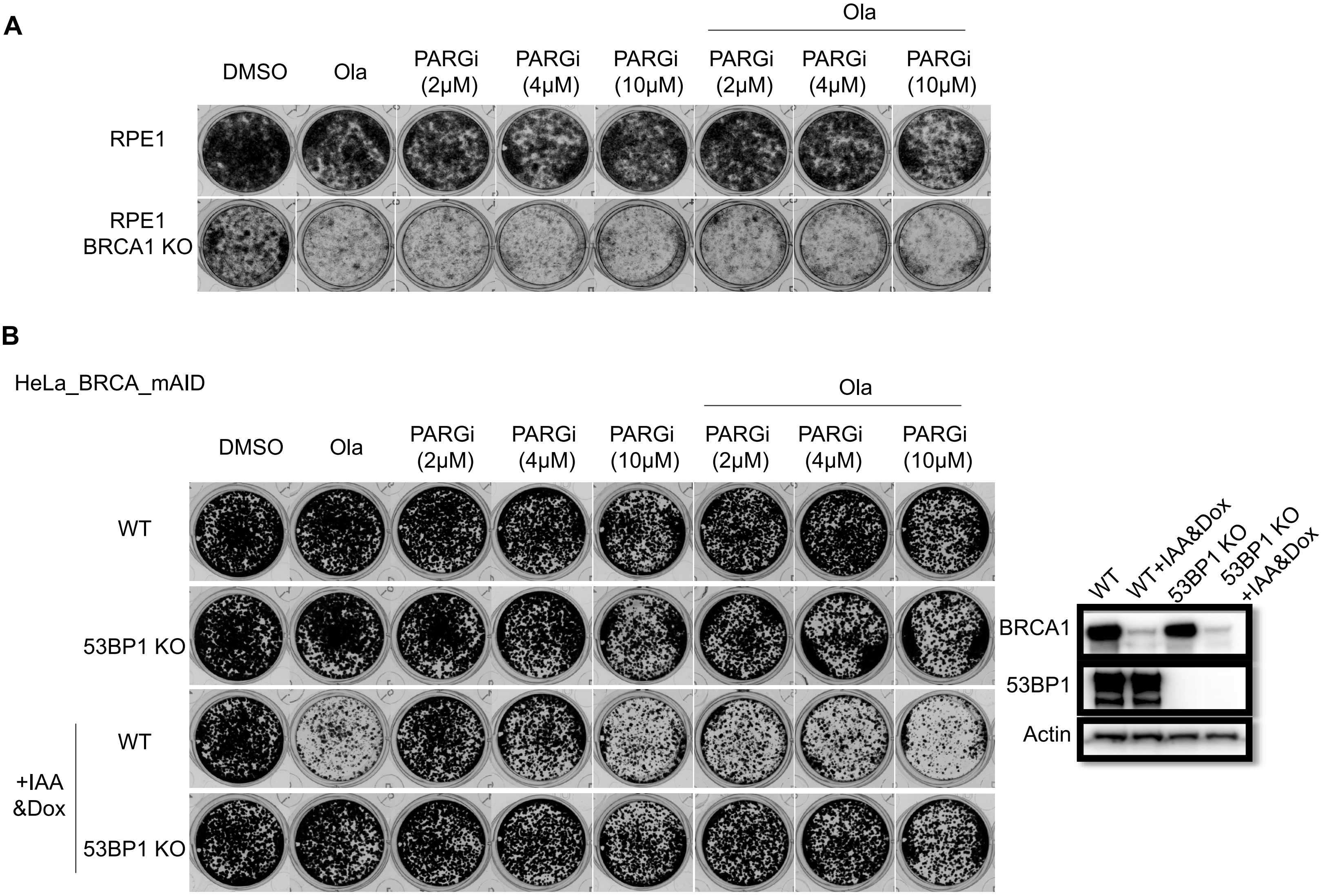

### Figure 6-Figure Supplement 2

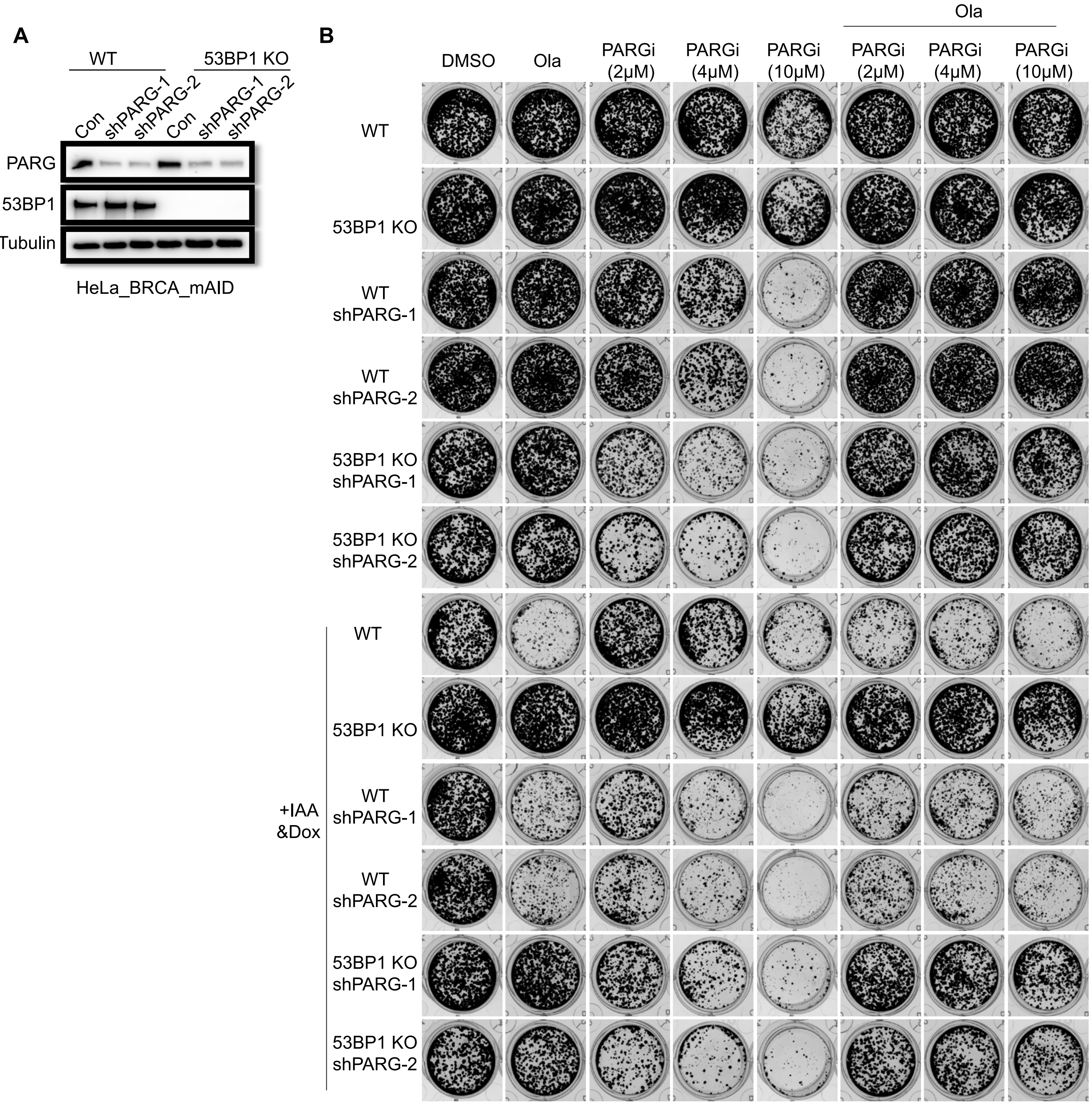

### Figure 7-Figure Supplement 2

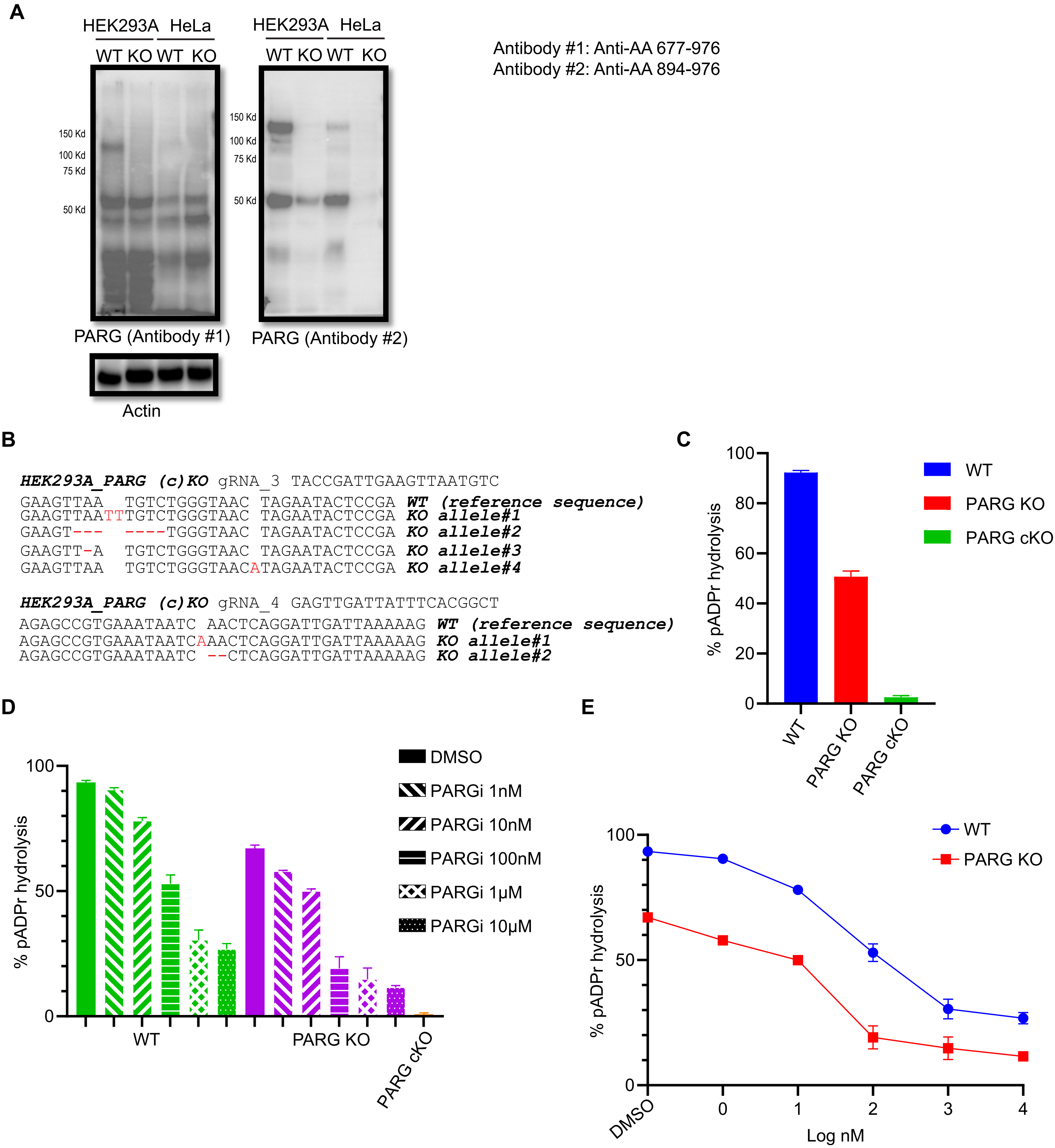
