## Supplementary material for "DePARylation is critical for S phase progression and cell survival": Figure 7-Figure Supplement 1

***HEK293A_PARG KO*** gRNA AGACTGCAGCAATGTGTAAG

AATGTGT AAGTGGCAAAATGAAGGGAAACACACGGAGCAGCTTTTGGAAAGTGAACCTCAAACAGTAA ***WT (reference sequence)***
AATGTGT ------------------------------------------------------ACAGTAA ***KO allele#1***
AATGTGT -----------TGAAGGGAAACACACGGAGCAGCTTTTGGAAAGTGAACCTCAAACAGTAA ***KO allele#2***
AATGTGTTAAGTGGCAAAATGAAGGGAAACACACGGAGCAGCTTTTGGAAAGTGAACCTCAAACAGTAA ***KO allele#3***

***HeLa_PARG KO*** gRNA TGCTATTCTGAAATACAATG

TTTTTTTGTTTCTGCATTTTCCACAGGATGCTATTCTGAAATACAA TGTGGCATATTCTA ***WT (reference sequence)***
TTTT---GTTTCTGCATTTTCCACAGGATGCTATTCTGAAATACAC TGTGGCATATTCTA ***KO allele#1***
TTTTTTTGTTTCTGCATTTTCCACAGGATGCTATTCTGAAA----A TGTGGCATATTCTA ***KO allele#2***
TTTTTTTGTTTCTGCATTTTCCACAGGATGCTATTCTGAAATACAAATGTGGCATATTCTA ***KO allele#3***

***HEK293A_PARG/PARP1/2 TKO*** PARP1_gRNA ATGGGTTCTCTGAGCTTCGG

TTGAGGTGGATGGGTTCTCTGAGCTT CGGTGGGATGACCAGC ***WT (reference sequence)***
TTGAGGTGGATGGGTTCTCTGAGCTTTCGGTGGGATGACCAGC ***KO allele#1***
TTGAGGTGGATGGGTTCTCTGAGC--+CGGTGGGATGACCAGC ***KO allele#2 (del 2bp and insert 112bp)***

***HEK293A_PARG/PARP1/2 TKO*** PARP2_gRNA ACAGCTAGATCTTCGGGTAC

TAGAGTCACAGCTAGATCTTCGGGTACAGGAG ***WT (reference sequence)***
TAGAGTCACAGCTAGATCTTC-GGTACAGGAG ***KO allele***

***HEK293A_POLB KO#2*** gRNA CCGCAGGAGACTCTCAACGG

TTCCCCCGTTGAGAGTCTCCTGCGGCGCCTTC ***WT (reference sequence)***
TTCCCC----GAGAGTCTCCTGCGGCGCCTTC ***KO allele#1***
TT-------TGAGAGTCTCCTGCGGCGCCTTC ***KO allele#2***

***HEK293A_POLB KO#4*** gRNA CAAGTACAATGCTTACAGGT

CTGCACTGTCCCACCT GTAAGCATTGTA ***WT (reference sequence)***
CTGCACTGTCCCACCTTGTAAGCATTGTA ***KO allele***

***HEK293A_XRCC1 KO#3*** gRNA CTACAGCAAGGTACCTAGGG

CCGGGTCAAAATTGTTTGCAGCCAGCCCTACAGCAAGGTACCTAGGGTGGGAGCCAG ***WT (reference sequence)***
CCGGGTCAAAATTGTTTGCAGCCAG----------------------TGGGAGCCAG ***KO allele#1***
CCGGGAG-------------------------------------GGGTGGGAGCCAG ***KO allele#2***

***HEK293A_XRCC1 KO#4*** gRNA GTATCGGCCAGACTGGACCC

CTGTCCCGGGT CCAGTCTGGCCGATACTTGG ***WT (reference sequence)***
CTGTCCC--GT CCAGTCTGGCCGATACTTGG ***KO allele#1***
CTGTCCCGGGTTCCAGTCTGGCCGATACTTGG ***KO allele#2***

***HEK293A_LIG1 KO#1*** gRNA TCACGGACTCGAATAAACCG

TCACGGACTCGAATAAACCGAGGGAAGC ***WT (reference sequence)***
TCACGGACTCGAAT----------AAGC ***KO allele***

***HEK293A_LIG1 KO#4*** gRNA CAAGAACAACTATCATCCCG

CATCTTCCACGGGATGAT AGTTGTTCTTGGCAG ***WT (reference sequence)***
CATCTTCCAC-GGATGAT AGTTGTTCTTGGCAG ***KO allele#1***
CAT---------------+AGTTGTTCTTGGCAG ***KO allele#2 (del 15bp and insert 53bp)***

***HEK293A_ARH3 KO#1*** gRNA GTGTAGTACAAGGCTTCTGT

GTGTAGTACAAGGCTTCT GTGGGGAGAGGA ***WT (reference sequence)***
GTGTAGTACAAGGCTTCTTGTGGGGAGAGGA ***KO allele#1***
GTGTAGTACAAGG----T GTGGGGAGAGGA ***KO allele#2***

***HEK293A_ARH3 KO#2*** gRNA GTGTAGTACAAGGCTTCTGT

TCCCTCCTCTCCCCACA GAAGCCTTGTAC ***WT (reference sequence)***
TCCCTCCTCTCCCCACA+GAAGCCTTGTAC ***KO allele (insert 118bp)***

***HEK293A_PARG/ARH3 DKO#1*** gRNA GTGTAGTACAAGGCTTCTGT

ATCTGTGTAGTACAAGGCTTCTGTGGGGA ***WT (reference sequence)***
ATCTGTGT----------------GGGGA ***KO allele#1***
ATCTGTGTAGTAC--------TGTGGGGA ***KO allele#2***

***HEK293A_PARG/ARH3 DKO#2*** gRNA GTGTAGTACAAGGCTTCTGT

GTGTAGTACAAGGCTTC TGTGGGGA ***WT (reference sequence)***
GTGTAGTACAAGGCTTCATGTGGGGA ***KO allele***

***HEK293A_HPF1 KO#3*** gRNA GACCAACTAATTGAAGTCCA

ATCATAAGGACCAACTAATTGAAGTCCAAG GCTTGCAGAAAG ***WT (reference sequence)***
ATCAT----------------------AAG GCTTGCAGAAAG ***KO allele#1***
ATCATAAGGACCAACTAATTGAAG------+GCTTGCAGAAAG ***KO allele#2 (del 6bp and insert 286bp)***

***HEK293A_HPF1 KO#2*** gRNA GACCAACTAATTGAAGTCCA

AATTGAAGTCCAAGGCTTGCAGAAAG ***WT (reference sequence)***
AATTGAAGT---AGGCTTGCAGAAAG ***KO allele#1***
AAT-----------------AGAAAG ***KO allele#2***

***HEK293A_PARG/HPF1 DKO#2*** gRNA GACCAACTAATTGAAGTCCA

GACCAACTAATTGAAG T CCAAGGCTTG ***WT (reference sequence)***
GACCAACTAATTGAAGTT CCAAGGCTTG ***KO allele#1***
GACCAACTAATTGAAG TCCCAAGGCTTG ***KO allele#2***

***HEK293A_PARG/HPF1 DKO#3*** gRNA GACCAACTAATTGAAGTCCA

GACCAACTAATTGAAG TCCAAGGCTTG ***WT (reference sequence)***
GACCAACTAATTGAAGTTCCAAGGCTTG ***KO allele#1***
GACCAACTAATTGAAA -CCAAGGCTTG ***KO allele#2***
